# A unified haplotype-based method for accurate and comprehensive variant calling

**DOI:** 10.1101/456103

**Authors:** Daniel P Cooke, David C Wedge, Gerton Lunter

## Abstract

Haplotype-based variant callers, which consider physical linkage between variant sites, are currently among the best tools for germline variation discovery and genotyping from short-read sequencing data. However, almost all such tools were designed specifically for detecting common germline variation in diploid populations, and give sub-optimal results in other scenarios. Here we present Octopus, a versatile haplotype-based variant caller that uses a polymorphic Bayesian genotyping model capable of modeling sequencing data from a range of experimental designs within a unified haplotype-aware framework. We show that Octopus accurately calls *de novo* mutations in parent-offspring trios and germline variants in individuals, including SNVs, indels, and small complex replacements such as microinversions. In addition, using a carefully designed synthetic-tumour data set derived from clean sequencing data from a sample with known germline haplotypes, and observed mutations in large cohort of tumour samples, we show that Octopus accurately characterizes germline and somatic variation in tumours, both with and without a paired normal sample. Sequencing reads and prior information are combined to phase called genotypes of arbitrary ploidy, including those with somatic mutations. Octopus also outputs realigned evidence BAMs to aid validation and interpretation.

---

Haplotype-based approaches have emerged as the method of choice for calling germline variants because these methods are robust to alignment errors from read mappers and have better signal-to-noise characteristics than positional approaches^1–7^. However, existing haplotype-based variant callers have several limitations. First, existing tools are sub-optimal for many problems as most implement models that assume either diploidy^1–3^ or constant copy number^4–6^, and assume that samples are selected from an idealized population of unrelated individuals. Such models are appropriate for calling germline variants in small cohorts, but provide a poorer fit to data generated in other experimental designs, such as studies involving samples with known relatedness such as paired tumours, single-cells, and parent-offspring trios, or in pooled tumour and bacterial sequencing where samples are often heterogeneous. These limitations cause researchers to implement custom pipelines that may integrate various callers and involve *post hoc* filtering and interpretation^8–16^. Second, existing haplotype-based methods suffer from windowing artifacts as variants are evaluated in independent non-overlapping regions. This can lead to false calls in complex regions where reads support variants that fall outside the region being evaluated. Third, existing methods do not make a clear distinction between the haplotype sequence supported by the read data, and the mutation events that gave rise to it. This makes it challenging to assign appropriate prior probabilities to these haplotype sequences, because different sets of mutations can have very different biological plausibility, despite giving rise to the same haplotype sequence. Fourth, haplotype-based methods, by nature, are able to physically phase variants, but existing tools are limited to phasing diploid genotypes, and none report potentially clinically relevant ^17^ phase information for somatic mutations with respect to germline variants or other somatic mutations.

To meet the growing demand for variant calling in divergent experimental designs, we designed an algorithm that can accommodate distinct genotype models within a unified haplotype-aware framework. We took inspiration from particle filtering ^18^ and developed a novel haplotype inference procedure that typically produces longer haplotypes than other methods, reducing the chance of windowing artifacts and improving the signal-to-noise ratio, resulting in more accurate variant calls. Furthermore, our method can propose and compare haplotypes composed of distinct sets of mutation events that nevertheless result in identical sequence, allowing us to consider the biological plausibility of mutations. We propose a probabilistic phasing algorithm that leverages both prior and read information, and can phase genotypes with arbitrary ploidy, including those that contain somatic mutations.

We present an implementation of our algorithm, *Octopus,* written in C++. We show that Octopus is more accurate than specialized state-of-the-art tools on several common experimental designs: germline calling in individuals; *de novo* calling in parent-offspring trios; and somatic variant calling in tumours, with and without paired normal samples. Octopus is freely available under the MIT license at https://github.com/luntergroup/octopus.

## RESULTS

### A unified variant calling algorithm

Octopus accepts sequencing data in the BAM and CRAM formats, and performs internal pre-processing, including PCR duplicate removal and adapter masking. Candidate variants are identified from the reads using a combination of local reassembly and pileup inspection with repeat awareness. In addition, variants from existing VCF files may also be considered. Haplotypes are then constructed exhaustively using a tree data structure (with nodes representing alleles and root-to-tip paths representing haplotypes) that is dynamically pruned, extended, and collapsed based on partial read evidence (**Fig. 1**). Calls are made once there is sufficient confident that haplotypes represented in the haplotype-tree explain all surrounding reads sufficiently well. Haplotype likelihoods are computed for each read and haplotype using a hidden Markov model (HMM) with context aware single nucleotide variant (SNV) and indel penalties. These likelihoods are the input to a polymorphic genotype *calling model,* the form of which depends on the experiment that generated the sequencing data **(Table 1**). Although calling models are responsible for calling variants, genotypes, and any other model specific inferences, each must be able to compute posterior distributions over haplotypes and genotypes, which are used for updating the haplotype-tree and for phasing, respectively. Variants are phased by evaluating the entropy of the computed genotype posterior distribution. Variant calls are then filtered, either with hard filters or a random forest classifier. Octopus optionally creates realigned evidence BAMs after calling by assigning and re-aligning reads to called haplotypes.

**Figure 1.**
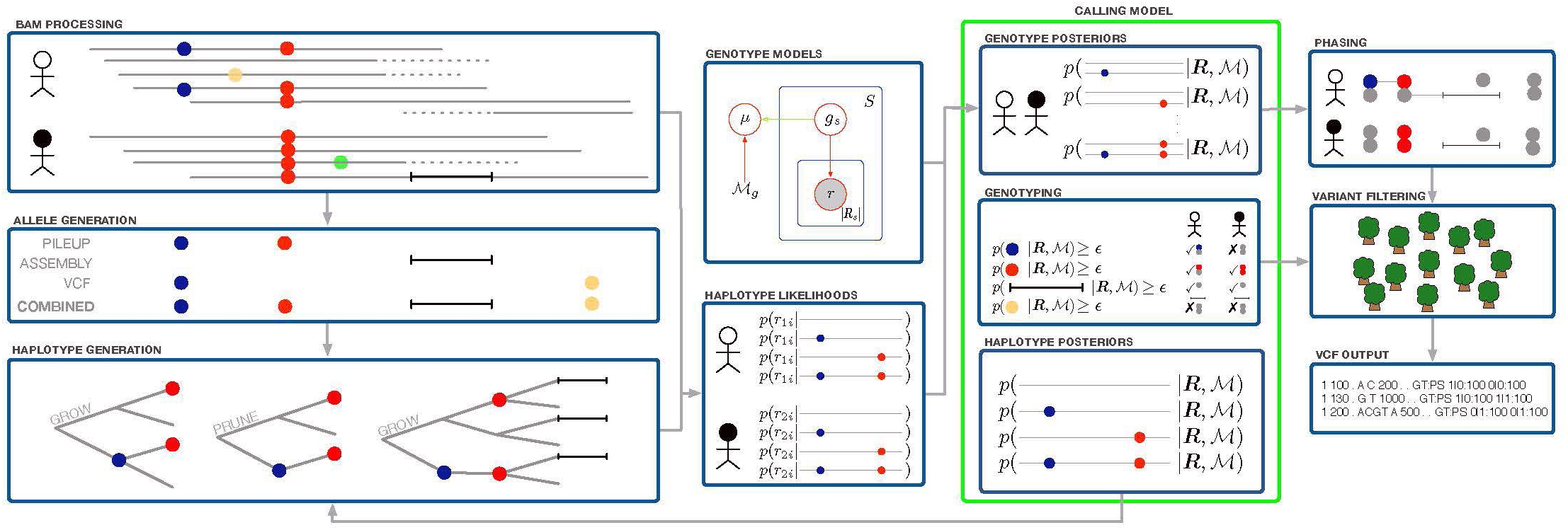
Overview of the unified haplotype-based algorithm, showing joint calling of two samples with the population calling model. Two SNVs (blue and red) are detected from read pileups, a deletion from local re-assembly, and a third SNV (yellow) from input VCF. The first two SNVs are added to the haplotype-tree, which then contains four haplotypes. After computing likelihoods for read-haplotype pairs, the haplotype posterior distribution computed by the calling model is used to prune the haplotype-tree by removing one haplotype (containing just the blue SNV). Next, the haplotype-tree is extended with the deletion, and the process repeats. The polymorphic calling model is shown in the green box. Only the population genotype model (Online Methods) is shown, in plate notation. Calling models also compute any model-specific inferences, such as *de novo* or somatic classification.

**Table 1.** Description of Bayesian calling models

| Model name | Class | Description |
| --- | --- | --- |
| Individual* | Germline | Calls germline variation in an individual with known ploidy. Haplotypes are expected to be observed at a frequency proportional to their copy number. |
| Population | Germline | Jointly calls germline variation in two or more samples with known ploidy but unknown relationship. Uses a joint genotype prior that can improve power to detect variation compared with individual calling by sharing information between samples. |
| Trio* | Germline<br>De novo | Jointly calls inherited and *de novo* germline variation in a diploid parent-offspring trio. By explicitly modelling inheritance patterns and *de novo* mutations, the model has higher power compared with typical joint calling. Allosome calling is supported. |
| Cancer* | Germline<br>Somatic | Jointly calls germline and somatic variation in paired tumour-normal or tumour only samples. The number of somatic haplotypes and their frequency are inferred from the data and used to call variants. Multiple tumours from the same individual may be called jointly. |
| Polyclone | Germline | Calls variation in an unknown mixture of haploid clones. Such samples often arise in bacterial or viral sequencing data where multiple clones may form due to sample contamination, mixed infection, or in-host evolution. The number of haplotypes and their frequency are inferred from the data. |

\* Evaluated in this article.

### Germline variants in individuals

To assess germline calling accuracy, we called variants in three well-characterized Genome in a Bottle (GIAB)^19^ samples: HG001 (NA12878), HG002 (NA24385), and HG005 (NA24631), in addition to the synthetic-diploid (Syndip) sample CHM1-CHM13^20^ that includes a validation set compiled using an approach orthogonal to that used for the GIAB truth sets. To account for different sequencing conditions we tested several HG001 and HG002 replicates, including two libraries prepared using the 10X Genomics Chromium protocol **(Supplementary Note 1**). We downloaded publicly available BAM files for the two 10X libraries (from GIAB) and for CHM1-CHM13 (from the Broad institute). BWA-MEM ^21^ was used to map all other data from raw FASTQ files. We compared Octopus to GATK4 ^6^, DeepVariant ^3^, Strelka2 ^2^, FreeBayes^5^, and Platypus^1^. We ran each caller according to the authors’ recommended settings **(Supplementary Note 2**). We trained Octopus’ random forest classifier using three independent NA12878 replicates **(Supplementary Note 1**) to filter variants. Default filters were used for DeepVariant and Strelka2. For Free-bayes and Platypus, we tried recommended hard filters, but found that this deteriorated performance as quantified by the F-measure (the harmonic mean of precision and recall), so we did not apply filters other than those based on variant or genotype quality (QUAL and GQ). Similarly, we found that using VQSR filtering for GATK4 - as recommended by the best practice guidelines - degraded performance, so we only used QUAL for filtering GATK4 calls. All calls were evaluated with RTG tools vcfeval ^22^.

Octopus had the highest F-measure on all tests other than the HG005 test **(Fig. 2b** and **Supplementary Table 1**). Performance differences were marginal between Octopus, Deep-Variant, and Strelka2 on the two Precision FDA Truth tests and GIAB HG005 test, all of which use data from GIAB sequenced on the Illumina HiSeq 2500 platform. Octopus substantially outperforms other callers on the two 10X Genomics samples, which have lower coverage and shorter read lengths than the other samples **(Supplementary Note 1**), in addition to barcoded library preparation. It is unclear why Strelka2 does not perform well on these data. Octopus also had considerably better performance on the Precision FDA Consistency and Syndip tests, both of which use data from the Illumina HiSeq X Ten platform, but were sequenced in different laboratories (Garvan institute and Broad institute, respectively). The HiSeq X Ten data had higher error rates than the other data, suggesting that Octopus is more robust to noise than other methods.

**Figure 2.**
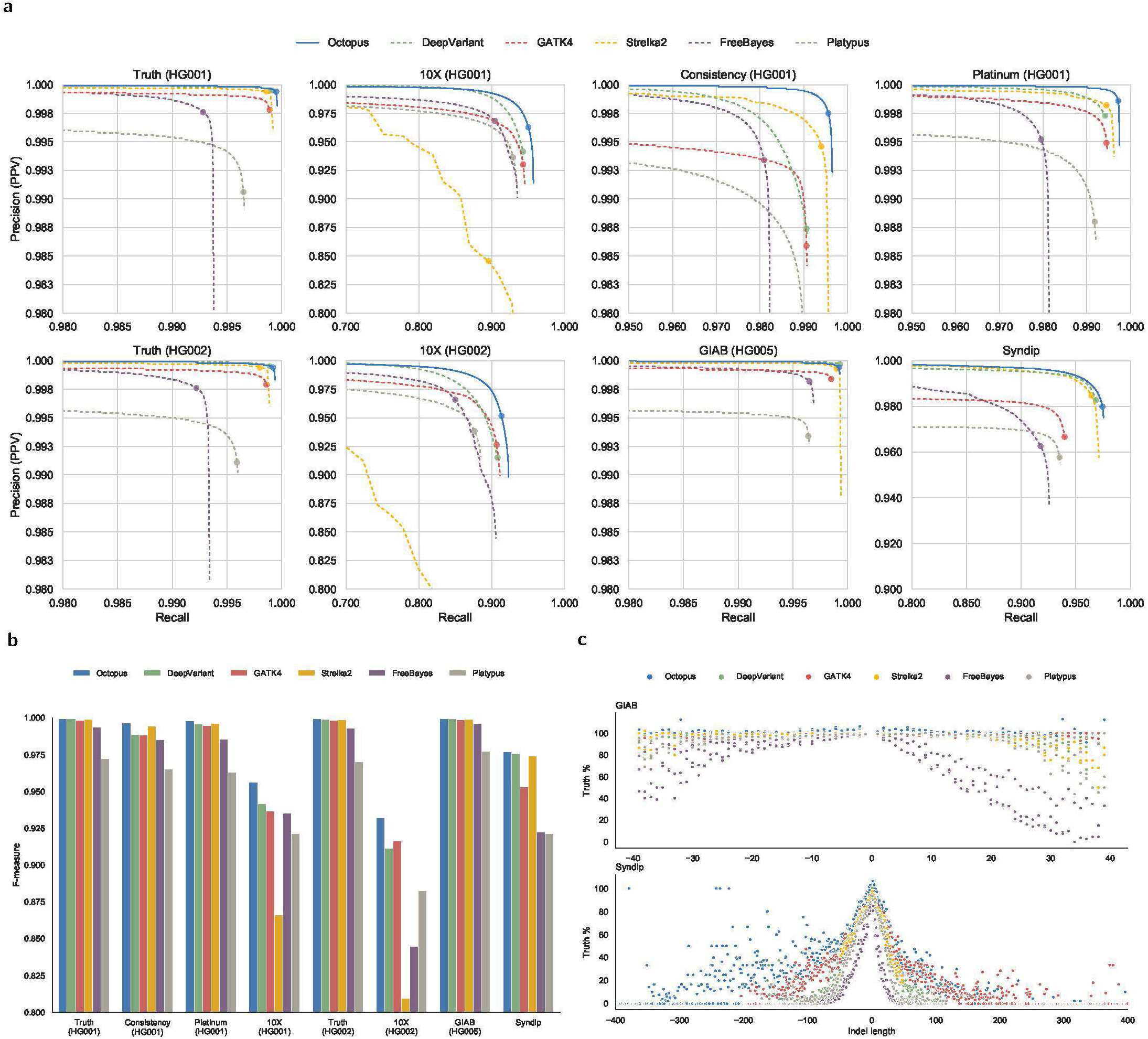
Germline variant calling accuracy. Comparison of Octopus with other methods on PrecisionFDA Truth Challenge HG001, GIAB 10X HG001, PrecisionFDA Consistency Challenge Garvan, Platinum Genomes HG001, PrecisionFDA Truth Challenge HG002, GIAB 10X HG002, GIAB HG005, and Syndip (CHM1-CHM13). The average sequencing depths of each dataset are approximately 50x, 34x, 40x, 50x, 50x, 25x, 50x, and 45x, respectively. All comparisons to the GIAB (version 3.3.2) and CHM1-CHM13 (version 0.5) truth sets were performed using RTG Tools vcfeval (version 3.9.1). **a** Precision-recall curves showing accuracy on all test sets. Scoring metrics used to generate curves were RFQUAL (Octopus), GQ (DeepVariant), QUAL (GATK4), GQX (Strelka2), GQ (FreeBayes), and QUAL (Platypus). The dots show typical PASS thresholds: 3 for Octopus, DeepVariant, and Strelka2; 20 for GATK4, FreeBayes, and Platypus. **b** F-Measures at PASS thresholds for each test set. **c** Proportions of true indels called in comparison to the number in the truth set by indel length. Positive lengths are insertions; negative lengths are deletions. Top: GIAB HiSeq tests (PrecisionFDA Truth Challenge HG001 & HG002, and GIAB HG005). Bottom: Syndip. The Syndip validation set has a larger range of indel sizes than the GIAB validation sets.

Contrary to common practice, we did not stratify our evaluation into SNVs and indels, because the true mutations that results in a haplotype are generally uncertain. For example, a sequence change of …AAACCC… to …AACCCC… could be explained either by a single SNV, or two homopolymer indels. We found that the representation used for the ground truth is biased towards the tools used to derive it. To demonstrate this, we compared the proportion of indels classified as true on the basis of haplotype matches, with the number of indels in the respective truth sets for the two Precision FDA Truth tests and GIAB HG005 tests. We observed that Octopus had on average 1.5% more ‘true’ indels than the total number of indels in the validation sets, while all other callers called at most 0.5% fewer true indels than in the validation set **(Fig. 2c),** despite there being less than 0.1% difference in overall sensitivity between Octopus and DeepVariant on these three tests. This apparent contradiction is due to Octopus having made indel calls in regions where other tools - and the validation sets - call SNVs, that result in the same haplotype sequence. We observed similar behaviour in the Syndip test for indel lengths up to 5bp **(Fig. 2c)**.

### Microinversion detection

Octopus calls complex replacements when an observed sequence cannot be satisfactorily explained in terms of simple SNVs and indels. Microinversions are one such complex replacement that involve the inversion of small (e.g. 3 – 100bp) tracts of DNA. Microinversions have not been comprehensively studied, likely due to calling difficulties, but have been suggested to play a role in evolution ^23^ and disease, including cancer ^24^. A method developed to identify microinversions reported an average of 3.8 per individual in 1000G data ^24^. We identifed a total of 152 microinversions ranging from 3 to 98 bp in the four NA12878 high-coverage replicates, of which 104 were called in at least three of the replicates and 75 were called in all four. We also found 103 microinversions in the NA24385 Precision FDA Truth sample, including a 3 bp inversion in a coding region of TTC6. The majority of called microinversions appear polymorphic; 72 of those detected in NA24385 were also called in NA12878.

Some of the microinversions called by Octopus have dbSNP entries. However, most also have decomposed entries (SNVs and indels) that are more commonly reported. For example, a 3 bp microinversion called by Octopus in NA12878 in a splice site of LIMD1 - a possible tumour suppressor - has a dbSNP entry (rs71615396). However, there are also entries for 3 SNVs (rs62242177, rs62242178, rs63132361) composing the microinversion. The SNV entries have frequency meta-information while the microinversion does not. Similarly, a 27 bp microinvserion in the 3’-UTR of PREPL, a gene linked with congenital myasthenic syndrome, has a dnSNP entry (rs71416108) without frequency information, but also has several decomposed entries, including SNVs and indels, supporting conflicting representations of the microinversion (e.g. rs368481611, rs373223610, rs1281267686), with frequency information.

### *De novo* mutations in parent-offspring trios

Random germline *de novo* mutations resulting from imperfections in the DNA replication process during meiosis provide the necessary genetic variation for evolution, and are causative of several Mendelian and polygenic diseases ^25–27^. The number of *de novo* mutations per genome duplication event is estimated to average around 70 per meiosis in humans ^28^. However, there is uncertainty in this estimate because accurate calling of *de novo* mutations remains challenging ^12–15,28^.

To assess *de novo* calling performance, we ran Octopus using the trio calling model on whole-genome data from a previously characterized WGS500 parent-offspring trio ^1^. We selected these data as the libraries were prepared directly from blood rather than cell lines. We compared calls made by all other tools evaluated for the germline analysis, set up for joint calling **(Supplementary Note 2).** In addition to the 63 *de novo* mutations previously Sanger validated in this sample, we manually identified a further 33 mutations by inspecting unfltered *de novo* calls made by three or more callers, as well as all passing *de novo* calls from Octopus and GATK4, using realigned BAMs that both GATK4 and Octopus are able to generate **(Supplementary Fig. 1**).

Only Octopus and GATK4 called a plausible number of *de novo* mutations **(Table 2**). Platypus and FreeBayes called approximately 2*x* and 3*x* more *de novo* mutations than the maximum expected. DeepVariant and Strelka2, despite being the two most accurate germline callers behind Octopus, called considerably more false positive *de novo* mutations than the other tools, demonstrating that strong germline calling performance does not guarantee accurate *de novo* calls. While we are confident the performance of the other callers could be improved with additional filtering, it is not always obvious how this is best achieved. For example, filtering DeepVariant calls by GQ resulted in almost complete loss of sensitivity before the number of false positives fell below 100 (GQ: 46, TP SNV: 9, TP INDEL: 0, FP: 90), while for Strelka2 we found that accuracy was highly sensitive to filtering by GQX (72 TP and 106 FP for GQX ≥ 22; 50 TP and 41 FP for GQX ≥ 23).

**Table 2.** *De novo* mutations called in WGS500 trio.

|  | TP SNVs | TP Indels | FN SNVs | FN Indels | FP |
| --- | --- | --- | --- | --- | --- |
| Octopus | 72 | 13 | 3 | 8 | 10 |
| DeepVariant | 70 | 0 | 5 | 21 | 10, 306 |
| Strelka2 | 75 | 5 | 0 | 16 | 1, 548 |
| GATK4 | 46 | 4 | 29 | 17 | 90 |
| FreeBayes | 73 | 5 | 2 | 16 | 339 |
| Platypus | 72 | 4 | 3 | 17 | 163 |

### Synthetic tumours

Comprehensive evaluation of somatic mutation calls is challenging because there is no gold standard reference material to compare with, and different tumour types have distinct mutation profiles ^29^. Although calls may be manually validated to obtain an estimate of the false positive rate, it is not straight-forward to estimate sensitivity as the ground truth remains uncertain. Although attempts have been made to accurately characterize somatic mutation profiles in real tumours by manual inspection ^30^, this process is limited by the sensitivity of existing tools, and is too time consuming to perform across a range of tumour types. An alternative strategy is to mix reads from unrelated individuals to create virtual tumours ^2,31^. However, this approach is unlikely to yield data with realistic mutation profiles, error profiles, and haplotype structure. A third approach is to spike mutations directly into raw sequencing reads from healthy tissue, which was the approach taken by the ICGC-TCGA-DREAM challenge ^32^.

We designed an unbiased and comprehensive somatic mutation calling performance test by improving the method used by the ICGC-TCGA-DREAM challenge to ensure that synthetic tumours would have realistic mutation profiles, error profiles, and haplotype structure **(Fig. 3**). We created two synthetic tumours by applying this method to reads from GIAB’s NA12878 high-coverage Illumina data **(Supplementary Note 3**). The first was derived from skin cancer mutations using a mutation rate of 1**/**kb (299, 873 mutations) and spike-in frequencies uniformly sampled between 2.5% and 50%, while the second was derived from breast cancer mutations using a mutation rate of 1**/**Mb (5, 956 mutations) and spike-in frequencies uniformly sampled between 0.5% and 20% **(Supplementary Fig. 2**). We used uniform spike-in frequencies, rather than simulating sub-clonal architecture, so that we could more thoroughly assess sensitivity across a range of variant allele frequencies. The average read depths were 60x and 65x for the synthetic skin and breast tumours, respectively. Independent normal samples were created from leftover reads with average depths of 30x and 35x for the synthetic skin and breast tumours, respectively.

**Figure 3.**
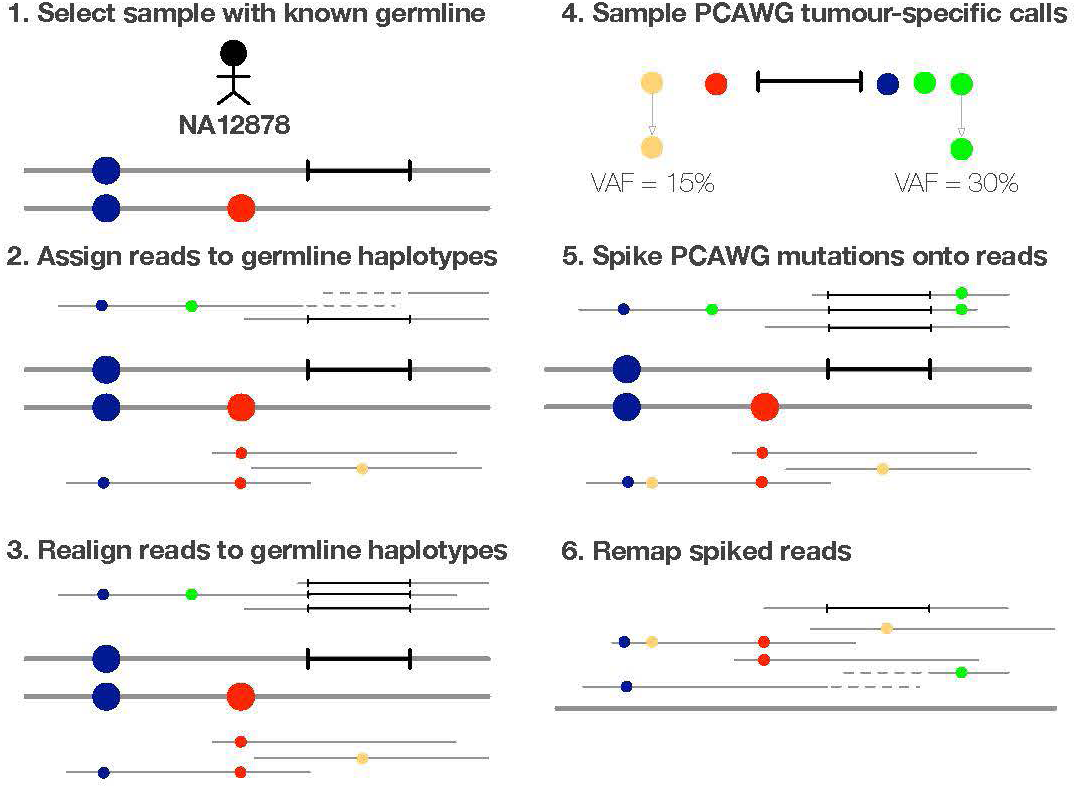
Overview of synthetic tumour creation. We used germline sequence data from a sample for which high-quality germline haplotypes are available (NA12878), and assigned and realigned reads to these haplotypes (Online Methods). This ensures that mutations are spiked onto consistent germline haplotypes and minimizes spike-in errors due to indels. We used spike-in mutations from tumour-specific whole-genome somatic mutation calls from the pan-cancer analysis of whole genomes (PCAWG) consortium ^33^ to ensure realistic somatic mutation profiles. Mutations were spiked in using a modified version of BAMSurgeon ^32^ (Online Methods). Reads were merged and remapped before variant calling to remove all realignment information.

### Somatic mutations in paired tumour-normal samples

We evaluated the accuracy of Octopus at calling somatic mutations in tumour-normal paired samples by calling variants in the skin and breast synthetic tumours. Calls were compared to Mutect2 ^31^, Strelka2 ^2^, LoFreq ^34^, Lancet ^35^, VarDict ^36^, and Platypus. We included Platypus, despite not being advertised as a somatic variant caller, to contrast germline and somatic callers, and because we are aware that germline callers (including Platypus) are sometimes integrated into somatic mutation calling pipelines. We trained Octopus’s random forest classifier on chromosome X of the skin synthetic tumour data, which we removed from the test set. However, we noted that Octopus was comparatively less reliant on filtering than other methods **(Supplementary Fig. 3**). Recommended filters were used for other methods **(Supplementary Note 2**).

During the course of evaluation we discovered a small number of mutations that were not in the truth sets but appeared real. This is not surprising since these data are derived from cell lines. To discount such cases, we identified calls not in the truth set but called by at least 3 of the 7 callers tested and ignored these calls during evaluation (skin: 843; breast: 788). In addition, we found a small fraction of true mutations (skin: 2, 766; breast: 1) that were incorrectly spiked in by BAMSurgeon, which we also ignored during evaluation.

There was a clear trade-off between recall (sensitivity) and precision (positive predictive value) between callers **(Fig. 4a).** Mutect2 and Strelka2 had similar F-Measures **(Supplementary Table 2**) on the synthetic skin test (0.9263 and 0.9251, respectively) despite Mutect2 showing higher recall; VarDict had highest recall on both tests, but also had lowest precision; Lancet had moderate precision and recall compared to other methods; LoFreq had near perfect precision, but only Platypus had lower recall. Octopus had higher recall than all callers other than VarDict, and only slightly lower precision than Strelka2. The number of false positives called by each caller is similar in both tests suggesting that each caller had unique biases, although it is possible that some of these false positive calls are genuine cell line artifacts. Overall, Octopus had substantially higher F-Measure on both tests than all other methods **(Supplementary Table 2).**

**Figure 4.**
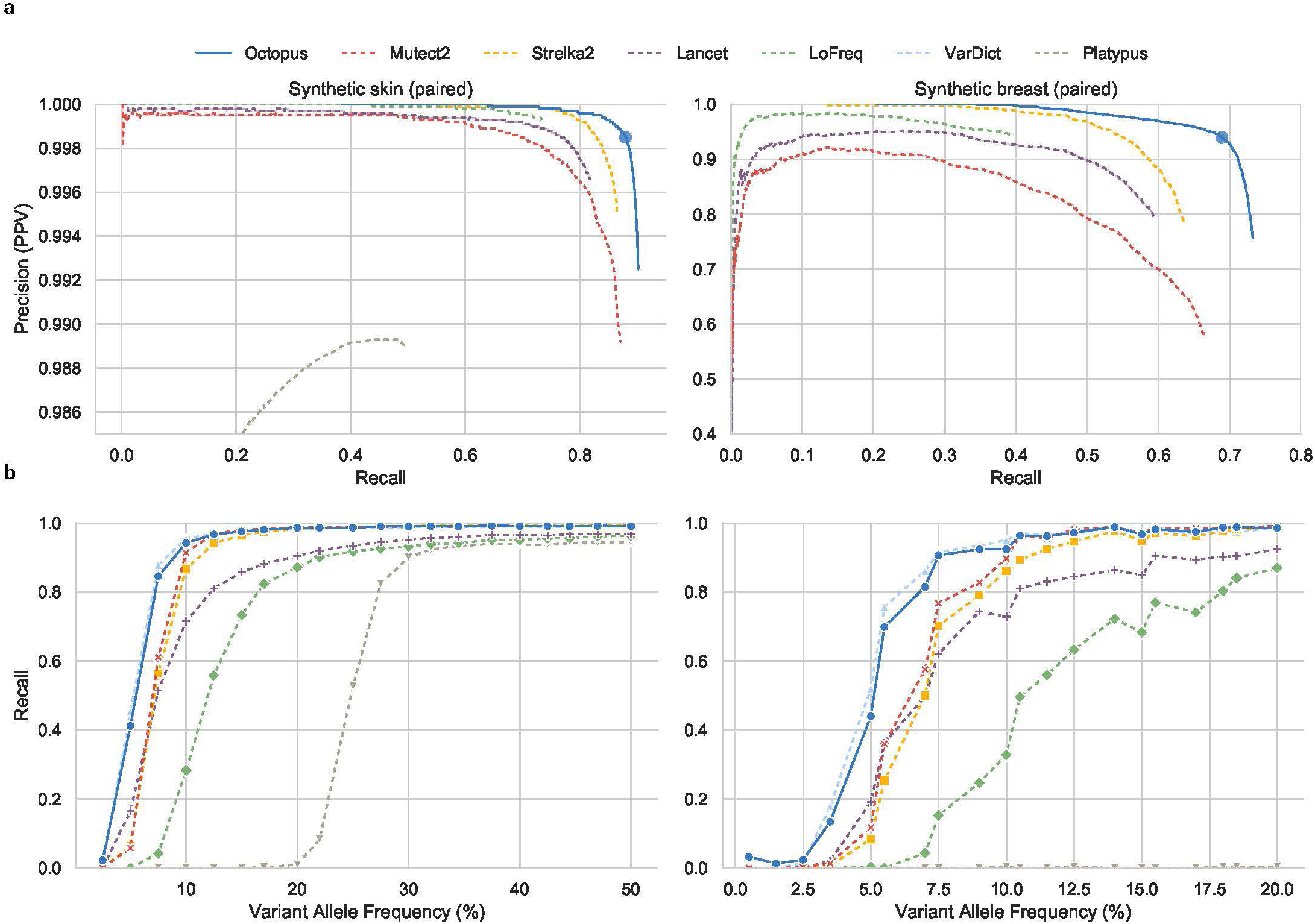
Somatic mutation calling accuracy for synthetic skin and breast tumours with a paired normal sample. **a** Precision-recall curves. Scoring metrics used to generate curves were RFQUAL (Octopus), TLOD (Mutect2), SomaticEVS (Strelka2), QUAL (Lancet), QUAL (LoFreq), SSF (VarDict), and QUAL (Platypus). Only PASS calls are used. VarDict is not visible as it is outside the axis limits due to low precision. Precisions on the two tests are substantially different as the skin set has almost 50 times as many true mutations as the breast set. Dots on the Octopus curve are placed at RFQUAL 7 (3 is used for the entire curve). **b** Recalls for each Variant Allele Frequency (VAF) using PASS variants. Points show true spike-in VAFs. The approximate depth for the synthetic skin and breast tumours were 60x and 65x, and 30x and 35x for their normal pairs, respectively. All comparisons to the synthetic tumour truth sets were performed using RTG Tools vcfeval.

Octopus also shows considerably better precision-recall trade-off than other callers, most notably at higher recalls. The call filter threshold for Octopus is set reasonably low by default (RFQUAL 3) to achieve high sensitivity, however, increasing this to 7 reduces the number of false positives by over a third in both tests, while only reducing the number of true positives by 2.5% and 6% in the synthetic skin and breast tests, respectively **(Fig. 4a).**

Most of the differences in recall, particularly between the best performing tools (Octopus, Strelka2, Mutect2, Lancet), were due to low frequency mutations **(Fig. 4b** and **Supplementary Fig. 5).** Sensitivity for mutations below 2.5% is poor for all callers (0.01 for Octopus and 0.002 for Mutect2). At 60x sequencing depth, a 2.5% VAF corresponds to an expectation of less than two observations. However, Octopus had considerably better sensitivity for mutations with VAFs between 4**−**10% (3-5 expected observations at 60x) and had only slightly worse recall than VarDict, which represents an approximate upper bound on sensitivity. Mutect2 had marginally better sensitivity at some moderate VAFs between 12.5% and 20%.

Finally, we re-ran all methods after downsampling the synthetic tumour and normal samples. We observed an even greater performance differential between Octopus and the other callers in all downsampled tests **(Supplementary Table 2** and **Supplementary Fig. 4-8).** In particular, Octopus was less affected by lower coverage in the normal sample than the other methods; Octopus had an F-Measure decrease of 0.9% in the synthetic skin test with half the normal depth, compared with 2.4% for Strelka2. Moreover, we found that Octopus had better overall performance calling somatic mutations in the synthetic skin tumour downsampled by 25% to 45x (F-Measure 0.9266) than all others methods did on the full 60x sample (maximum F-Measure 0.9263 from Mutect2).

### Mutations in tumour-only samples

Most somatic detection tools require a paired normal sample ^2,34,35^, but paired control tissues are not always available. We tested Octopus’s ability to call mutations in tumour-only data by calling variants in both synthetic tumours without providing the paired normal samples. We compared Octopus with a tool designed for calling unpaired tumour samples, Pisces ^37^.

Unsurprisingly, Octopus’ somatic calling accuracy is worse compared with the paired test, most notably the number of false positives is considerably higher on both tests. However, Octopus substantially outperforms Pisces both in terms of recall and precision **(Fig. 5a).** Pisces calls 6.5*x* and 8.3*x* more false positive somatic mutations than Octopus in the skin and breast tests respectively, yet Octopus calls 2.3*x* and 1.3*x* more true positives **(Supplementary Table 3).** The difference in sensitivity is primarily because Octopus is sensitive to allele frequencies between 5 and 30%, in contrast to Pisces, which is only sensitive to frequencies between 10 and 20% **(Fig. 5b).**

**Figure 5.**
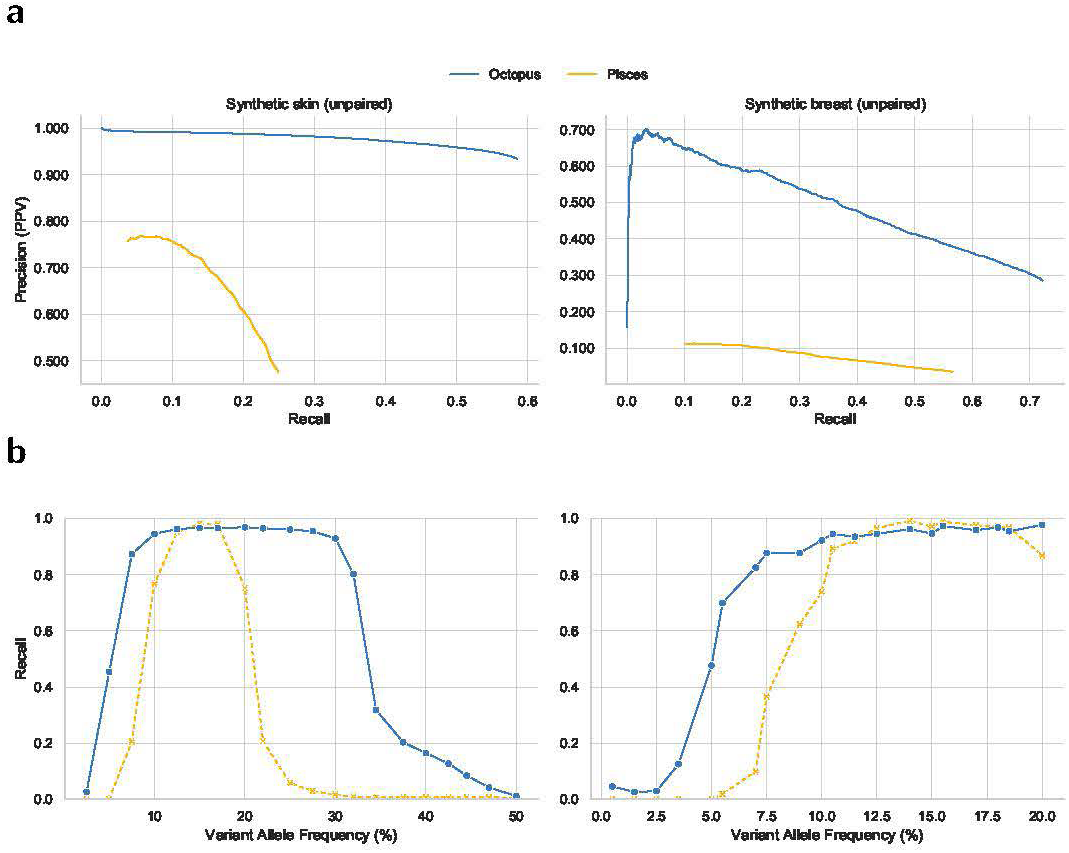
Somatic mutation calling accuracy in synthetic skin and breast tumours without a paired normal sample. **a** Precision-recall curves. Scoring metrics used to generate curves were RFQUAL (Octopus) and QUAL (Pisces). **b** Recalls for each Variant Allele Frequency (VAF). Germline and somatic calls were compared to the truth sets with RTG Tools vcfeval.

A significant challenge with tumour-only calling, compared to somatic calling with a paired normal sample (of reasonable depth), is correctly classification of variants as either somatic or germline. Both Octopus and Pisces are capable of calling and classifying germline and somatic variants, and we observed that both callers misclassified a large number of true somatic mutations as germline variants in the skin test, which largely explains the drop-off in recall at VAFs above 30% **(Fig. 5b).** Unlike Pisces, Octopus provides a measure of uncertainty in classification. Depending on the application, classification may or may not be important; it may be sufficient to know whether the variant is present or not. We therefore evaluated the performance of both callers on combined germline and somatic truth sets; ignoring somatic classification. We found that there was little difference in sensitivity between the callers, but a large difference in precision **(Supplementary Fig. 9).** Octopus called marginally more false positives on the combined test than on the somatic-only test (< 1, 700 in both tests), indicating that there are few germline calling errors, while Pisces calls over 17, 000 additional false positives in both tests.

### Phasing somatic mutations

In some situations, such as when compound heterozygous mutations are suspected, it is clinically relevant to be able to determine the germline haplotype affected by a mutation ^38^. Furthermore, phasing information is informative of tumour clonal architecture. To the best of our knowledge, no existing caller is able to phase somatic mutations, either with germline variants or other somatic mutations.

Since we designed the synthetic tumours used to benchmark somatic mutation calling so that individual reads would respect haplotype structure (but not necessarily read pairs), we know the local phase of all somatic mutations. We investigated Octopus’ ability to phase somatic mutations by evaluating how well phasing information was recovered in the paired synthetic skin tumour test. We found that of the 257, 930 somatic mutations that Octopus calls, 57, 217 (22%) were phased with one or more heterozygous germline variant. Furthermore, 9, 834 (4%) were phased with at-least one other somatic mutation, and 2, 848 (1% overall) of these were also phased with a heterozygous germline variant. We found that approximately 93% of reported somatic-germline phasings with phase quality 10 or greater (Phred scaled) were correct, indicating that the phase quality score is well calibrated.

## DISCUSSION

We have shown that Octopus is more accurate than state-ofthe art variant callers on several germline samples using two independent validation sets. Performance differences were most evident on the two 10X and two X Ten samples (Consistency and Syndip), arguably the most challenging tests because of lower read depths and higher sequencing error rates than in the other four samples. These results are likely because Octopus is better able to discern noise due to longer haplotypes, more effective error modelling, and more realistic mutation priors.

Our analysis of germline indels indicate that Octopus is able to call a wider range of indels than other methods. Octopus calls considerably more true short (< 15*bp*) indels than other methods, and even more than are represented in the truth sets. One explanation for this is that existing methods systematically miscall a large number of indels as SNVs in tandem repeat regions leading to under representation in the truth sets. Although both representations result in the same haplotype sequence, the distinction could have important clinical consequences, such as for mutation signature profiling^29^ or microsatellite instability analysis ^39,40^. Moreover, around half of our manually curated *de novo* calls occur in microsatellites. Such sites are known to have higher mutation rates than average but are almost always ignored in *de novo* mutation studies because such regions are also difficult to accurately call. Our results indicate that Octopus is sufficiently specific for these mutations to be considered. We also found that in the Syndip test, Octopus and GATK4 call considerably higher proportions of true large indels (> 50bp) than other callers, further supporting Octopus’ sensitivity for indels.

Octopus provides evidence that microinversions are more common than previously thought, suggesting that they warrant investigation for functional effects. In a follow-up experiment (data not shown), we found that Octopus called a considerable number of microinversions in *Mycobacterium tuberculosis* isolates, almost all of which were found within inverted repeat patterns, as is the case for many of the microinversions that we found in human genomes. Although we cannot rule out sequencing artifacts, inverted repeats are known to form cruciform extrusions causing genetic instability that results in high mutability, and have been shown to cause pathogenic mutations in cancer ^41^, and have previously been associated with inverted repeats in mammalian evolution ^23,42^.

High-throughput sequencing is widely used throughout the genomics community, yet the most powerful variant calling methods are optimized for human germline population data. A key advantage of Octopus is its polymorphic calling model, allowing it to optimize performance for other experimental designs as well, including for calling somatic variants from tumour samples.

Sensitivity to a wide range of variant allele frequencies is crucial for fully characterizing the mutation profiles of tumour genomes, however, our results suggest that existing somatic callers have poor sensitivity for variation occurring below 10% frequency at typical sequencing depths - unless a large number of false positives are accepted. Octopus shows near optimal sensitivity across all variant allele frequencies tested while remaining highly precise. Furthermore, Octopus also showed better precision-recall trade-off than other methods, showing that call sets can be refined on the basis of a single score. Given that only a single chromosome of data was used for training Octopus’ random forest used to score calls, it is likely that this aspect of performance could be improved even further by including additional training data. Octopus was more robust to changes in sequencing depth than other methods, both in the tumour and normal samples. In summary, Octopus out-performed the most accurate existing methods while using less than 85% of all the data.

Our analysis of somatic mutation phasing indicates that Octopus could be used to detect cases of bi-allelic loss-of-function mutations in tumours, and provide information on tumour clonal architecture - beyond the information already provided by variant allele frequency inference. Our phasing method works equally well for SNVs and indels and the algorithm only depends on genotype posterior distributions.

Octopus has a number of usability advantages over existing tools. For example, reads do not require pre-processing as this is done internally, simplifying workflows and eliminating the need for intermediate BAM files. As an example, we required over 20 commands to call *de novo* mutations using the GATK4 pipeline compared with a single command for Octopus. Furthermore, multithreading is built in, and disk access is optimized for fewer long accesses rather than many short accesses, improving I/O throughput. Octopus is capable of producing realigned ‘evidence’ BAMs, including for somatically mutated haplotypes. We hope that clinicians in particular will find this feature useful for aiding variant and phase call validation and interpretation.

Although we did not include an evaluation of the polyclone calling model, intended for bacterial or viral sequencing protocols, we are confident that Octopus would perform equally well for this type of data, since the statistical problems faced are similar to those found in tumour-only calling but without the complication of variant classification. We look forward to forthcoming validation sets ^43^ to verify this.

Overall, Octopus is highly accurate on several important experimental designs, demonstrating the advantage of our unified haplotype-based algorithm. As new technologies and experimental designs emerge, the flexibility of our method will allow us to rapidly incorporate new calling models that take full advantage of the information present in the data generated by each experiment.

## ONLINE METHODS

### Read pre-processing

Input reads are scanned in random sub-regions of the input region set to estimate basic statistics such as average depth and read lengths. These statistics are used to determine sub-regions of the input regions to buffer input reads, so that the memory occupied by read data is below a user-defined limit. If multiple threads are requested then the buffer limit is shared evenly between threads. Input read files can contain multiple samples, but must have associated read group information.

#### Transformations

Read transformations adjust the data contained in a read observation without removing the read. Most of the transformations re-calibrate base qualities in certain ways. By default, the only read transformations are to mask (set the base quality to zero) all bases considered to overlap sequencing adapters, and mask all but one base of bases overlapping other segments part of the same read template.

#### Filtering

Read filters remove reads that are likely problematic and cannot be transformed into something useful. Read filtering is applied *after* read transformations. Reads are removed if *any* of the given filtering predicates fails. By default, reads are filtered if they: i) Have malformed CIGAR strings. ii) Are unmapped. iii). Have mapping quality below 20. iv) Are marked as QC fails. v) Are identified as being duplicates by Octopus. vi) Are marked as duplicates. vii) Are secondary or supplementary alignments. viii) Have fewer than 20 bases with base quality 20 or greater.

#### Downsampling

Read downsampling removes reads to satisfy user-specified depth criteria. Sample read sets are downsampled independently. First, regions that have depth above a certain threshold are detected. Using input alignments, the number of bases that need to be removed are calculated for each position. Reads in are then removed iteratively by first selecting a position to downsample with probability proportional to the required depth reduction at each position, and then selecting a read overlapping that position with uniform probability. By default, regions with depth above 1, 000 are downsampled to reach an average read depth of 500.

### Candidate allele discovery

Candidate alleles are generated jointly for all samples by taking the union of candidates generated from a set of orthogonal methods (*generators*). Users can choose which generators to use to optimize accuracy and runtime. The read-backed generators can be tuned to increase sensitivity for low-frequency variation.

#### Pileups

Uses the read mapping and alignment information present in the input BAM files and proposes candidates bases on mismatches present in these alignments. Alleles are only proposed if the observation of a particular satisfies an inclusion predicate, which primarily depends on the observation frequency; observed base qualities; and observed read strands.

#### Local reassembly

Discards read alignment information (but keeps mapping location) and builds a kmer based assembly graph (i.e. *de Bruijn* graph) at regions considered likely to contain variation. Once the graph is constructed, paths with low observation kmer counts and cycles are pruned. Candidate alleles are extracted by enumerating the highest scoring non-reference bubbles, where the score is determined by the sum of the kmer counts, weighted by strand bias. The non-reference path of the bubble is aligned to the reference to form candidate alleles. Complex variation such as microinversions are found by inspecting bubbles.

#### Repeat realignment

Identifies common patterns of misalignments in tandem repeat regions that results in runs of regularly spaced SNV mismatches and proposes relevant indel candidates.

#### Input VCF

Reads a set of user-specified VCFs and extracts all alleles present in the input regions.

### Candidate haplotype generation

#### Haplotype-tree construction

Haplotypes are exhaustively constructed from all candidate allele combinations. This approach differs from other methods that construct haplotypes directly from read observations ^4,5^. The primary advantage of our method is that haplotypes with no direct read support are proposed, so the length of haplotypes is not limited by read length. However, since the number of haplotypes is exponential in the number of alleles this approach is usually only feasible for very short haplotypes. To allow construction of long haplotypes, we use a graph data structure, called a *haplotype-tree,* where tree-nodes are alleles and branches are haplotypes. The key property of the haplotype-tree is that individual branches (haplotypes) can be removed or extended, which allows us to limit the haplotypes in the tree to only the most likely ones given partial data.

#### Controlling tree growth and active regions

The size of the tree (number of haplotypes) is controlled by a user-defined parameter (default 200). The tree is grown by adding alleles sequentially in position order, until the size of the tree reaches the haplotype limit. Growth rate is also controlled by checking if reads overlapping alleles at the frontier of the tree overlap with alleles that are to be added next. Alleles that overlap with the tree frontier are always added, which can cause the tree to exceed the haplotype limit. In this case, the tree is either pruned (see filtering below) or some alleles (usually large deletions) are identified to be temporarily removed from the active set, and are added again later once the tree has been sufficiently pruned. The alleles in the tree are called *active.* Alleles that are active but have already been evaluated by the calling model are called *indicators.* Periodically, indicator alleles from the root of the tree are dropped to allow room for new active alleles. Only once alleles are dropped from the tree are they viable to be called. The frequency at which this is done is user-controlled, and may be turned off completely to give non-overlapping active regions. The default behaviour is to drop indicator alleles based on the current tree size, and by checking reads overlapping indicator alleles and novel active alleles.

#### Deduplication

It is possible that duplicate haplotypes (i.e. identical sequence) exist in the haplotype tree since candidate alleles are exhaustively combined. Duplicate haplotypes will have identical likelihoods as the probability of generating a read from a haplotype is only a function of the sequence itself. However, duplicate haplotypes may not have equal posterior probability as the prior probability of a haplotype segregating depends on the alleles which compose the haplotypes. Since the posterior of duplicate haplotypes is only dependent on the prior we just keep the duplicate with the greatest prior probability.

#### Filtering

There are two haplotype filtering stages: prior and post to genotype inference. The latter is always preferable as it is possible to deduce a marginal posterior probability for each haplotype segregating in the samples, which contains all available information. However, it is sometimes necessary to reduce the number of haplotypes considered by the genotype model as the haplotype-tree can exceed the provided haplotype limit. We considered a number of alternatives and found that likelihood based statistics are most effective. In particular, we rank haplotypes by the number of reads assigned to each haplotype calculated by maximum likelihood.

### Haplotype likelihood calculation

#### Remapping

The first step of the likelihood calculation is to remap reads to candidate haplotypes. This is required because the likelihood model requires that reads already be reasonably well placed, and the mapping position provided by the read mapper may not be accurate with respect to certain haplotypes (e.g. when indels are present).

We use a simple *k-mer* based mapper to find putative mapping locations. Briefly, the k-mer (*k* is hard coded) starting at each read and haplotype base are calculated. For each k-mer in the read we then check if the k-mer exists in the haplotype, and if so, calculate which position in the haplotype the read would start assuming perfect alignment between the read and haplotype up until the k-mer (i.e. offset by the k-mer position in the read). After doing this for all k-mers in the read we find positions in the haplotype that have high a high number of putative read starts and emit these mapping positions.

#### Error models

The pair HMM (described below) for calculating read likelihoods is parameterized by a sequencing error model with indel gap open and gap extension penalties, and optionally, SNV mismatch caps. The current implementation uses a constant gap extension penalty. The penalties are set according to local repeat context, up to some maximum repeat period (currently trinucleotide repeats). There are currently two sequence error models: the default, which is intended for typical Illumina HiSeq 2500 quality data, and one intended for platforms with higher error rates, such as the Illumina HiSeq X Ten. We did not use any automated inference procedure to arrive at the parameters for these two models, but set them based on experience and observation.

#### pHMM

Haplotype likelihoods are calculated using a pair hidden Markov Model (pHMM). The pHMM likelihood method computes the approximate Viterbi probability of the read given the haplotype. We use the Viterbi probability rather than the forward probability since the Viterbi probability is considerably cheaper to compute in log space, and in practice the difference between using the two probabilities is small. The simplest pHMM implementation has positional gap open penalties which are a parameter to the model. There is a second version of the pHMM that also has SNV mismatch caps: A vector of nucleotides and maximum base mismatch penalties - one for each position in the haplotype sequence - that limit the penalty of a mismatch aligned to that position where the mismatching read base is the nucleotide indicated in the vector at that position. The intention of this is to model a common error mode in sequencing data in repetitive regions, especially homopolymers, where a single base on the edge of the repeat ‘falls over’ to the leading base of the repeat.

The pHMM is a performance bottleneck in Octopus and it therefore uses a highly optimized banded SIMD implementation. Being banded, the likelihood calculation only explores parts of alignment space that results in indels less than the band size (currently 8). For short Illumina quality reads, this limitation is almost never an issue as indel *errors* greater than this are extremely rare. For longer, noisier reads, the band size would need to be increased. Since the band size is just the width of the SIMD register used by the implementation (currently SSE2), to achieve a larger band size we would need to modify the code to use SIMD instructions that support larger register sizes (e.g. AVX2).

#### Inactive flank scoring

If there are read observations that partially overlap the current set of active alleles, but also support inactive alleles, then the likelihood for a true haplotype could be lower than a false one only because the false one better supports a true haplotype that has yet to be considered. This, in short, is the ‘windowing’ problem that all haplotype methods must address. Octopus’ solution to the windowing problem is to only commit to calling candidate alleles once there is reasonable confidence that the reads supporting the alleles have had likelihoods evaluated on *all* the true alleles they support. However, the problem still remains that we must evaluate the likelihood function with a haplotype that is correct in the active region, but false outside this region (sequences outside the active region are always padded with reference). Our solution is to ‘discount’ any reductions to the likelihood that arise from mismatches outside the active region. We do this by retracing the Viterbi path and subtract terms from the log likelihood that are due to mismatches outside the active region.

#### Mapping qualities

Mapping quality is an important statistic that reflects the trustworthiness of a reads mapping location. Formally, it has been defined as the probability the read alignment is wrong. In Octopus, we are not as concerned with the *alignment* of an input read as with the mapping *location,* since all reads are realigned internally. If the read is incorrectly mapped to a degree that Octopus’ remapping step cannot place the read correctly, then the read is not informative of the true haplotype and should not be used. To account for mismapped reads, we optionally factor mapping quality into the read likelihood calculation using the formula:
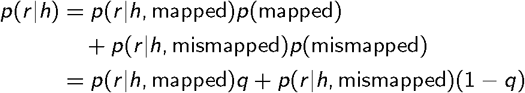
where 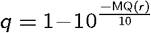. *p*(*r|h*, mapped) is then simply the likelihood from the pHMM. *p*(*r|h*, mismapped) is a more interesting quantity. If the read is unmapped, then it originated from some other sequence not localized to the reference region used to construct the haplotype under consideration. Since we cannot reasonably calculate this probability, we set it to a constant value (1). Although this assumption is not valid, the behaviour of the whole calculation to limit the impact of mismapped reads is achieved.

### Mutation models

#### Indel mutation model

Local gap open and extension probabilities are modelled for germline, *de novo*, and somatic mutations with a single indel mutation model. The model takes as input a base rate parameter which is scaled according to the local repeat composition of the sequence using the model in Montgomery et al. ^44^. Gap extension probabilities are assigned based on repeat composition and the current gap length. The extension model encourages the inclusion of whole repeat periods by assigning high probability to extensions of incomplete repeat periods, as indels in tandem repeats almost always occur in whole periods.

#### Coalescent mutation model

The coalescent mutation model, 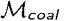, assigns probabilities to sets of haplotypes assumed to be sampled randomly from an idealized population. For a given set of haplotypes ***h*** = {*h*_1_, …, *h_m_*} we first calculate two the number of unique segregating SNV sites observed in ***h,*** *k*_1_, and the number of unique segregating indel sites observed in ***h,*** *k*_2_. Both *k*_1_ and *k*_2_ are calculated by comparing the alleles composing haplotypes to those in the reference haplotype.

The model has two parameters: the SNV heterozygosity, *θ*_1_, set depending on user input; and the indel heterozygosity, *θ*_2_, which is set according the the indel mutation model given the reference sequence and a user-supplied base indel heterozygosity. The maximum indel gap open heterozygosity for all segregating indels in ***h*** is used for *θ*_2_. Probability is then assigned to ***h*** by extending the distribution for the number of segregating sites under the coalescent model ^45^ with two heterozygosities:
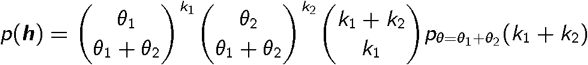
where 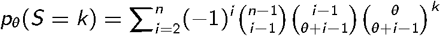.

We note that a limitation of this model is that only a single indel heterozygosity value is used for all positions in the observed haplotypes. While this assumption is unrealistic, it rarely detrimental since the most important aspect of the model is to assign high probability to indels in repeat regions. The likelihood of proposing some other spurious indel in a region outside the repeat region is small given the haplotype lengths normally considered.

#### De novo and somatic mutation models

The *de novo* mutation model is intended assign probabilities to *de novo* mutation occurring on a single haplotype during a single DNA replication. For haplotypes *h*_1_ and *h*_2_, the model assigns probabilities *p*(*h*_2_|*h*_1_) according to the indel mutation model and a SNV mutation rate that are parameters to the model. The somatic mutation model assigns probabilities to somatic mutations occurring on a single haplotype over some time period and is identical to the *de novo* mutation model.

### Genotype prior models

Genotype prior models are used to assign prior probability to arrangements of genotypes, *g* = (*h*_1_, …, *h_m_*) for ploidy *m.* There are two types of genotype prior models: single and joint. Single genotype prior models assign probability to single genotypes, 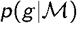, joint genotype prior models assign probability to a *combination* of genotypes, 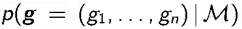. In some cases, such as the Coalescent genotype prior model, the former is simply a particular instance of the first. We report the unnormalized versions of each genotype prior model since normalization is trivial, and is always performed as part of the genotype posterior calculation.

#### Uniform genotype prior model

The uniform genotype prior model, 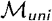, is the simplest genotype prior model. We have:
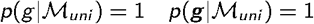
for the single case, and for the joint case, respectively.

#### HWE genotype prior model

The Hardy-Weinberg Equilibrium (HWE) genotype prior model, 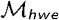, assigns probability to genotypes assuming HWE. The model is paramertized by a set of known haplotypes ***h*** = {*h*_1_, …, *h_m_*}, and their frequencies, *f_i_* (for *i* = 1 to *m*). The haplotype frequencies may be set explicitly, or calculated with empirical Bayes. The HWE prior is a multinomial distribution:
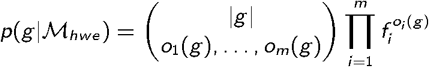
where *o_i_*(*g*) is the number of haplotype *i* occurrences in genotype *g.*

#### Coalescent-HWE genotype prior model

The coalescent-HWE genotype prior model, 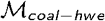, is suitable for modelling genotypes randomly sampled from an idealized population; it is the default germline prior model for all calling models when sample relationship is unknown. There are two components to this model: a *segregation* model, that assigns probability to the pattern of observed alleles in the genotype(s); and a *frequency* model that assigns probability to the frequency each haplotype is observed. In particular, the segregation model is just the coalescent mutation model and the frequency model is a Hardy-Weinberg model parameterized with empirical Bayes. We then have:
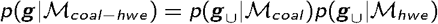
where *g*_∪_ = {*h* ∊ *g* : *g* ∊ *g*}, for the joint case. The individual case can be simplified to
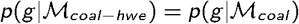
when |*g*| ≤ 2 (i.e. the sample is haploid or diploid) since 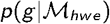 is then constant.

#### Trio genotype prior model

The trio genotype prior model, 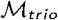, assigns probabilities to triplets of genotypes that originate from parent-offspring trios. This model encapsulates two elements of uncertainty: inheritance patterns, and parental haplotype modification due to *de novo* mutations. The model uses Coalescent-HWE genotype prior model or uniform prior model to assign probability to parental genotypes and the *de novo* mutation model to model modifications of parental haplotypes. Letting *g_m_, g_p_, g_o_* denote the maternal, paternal, and offspring genotypes, respectively. The model calculates
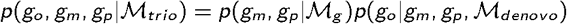

The form of the latter term 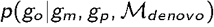 is dependent on meiosis and fertilization in the species under consideration. We only consider the mammalian case.

Writing 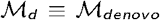 for brevity. In the autosomal (i.e. all diploid) case, we have
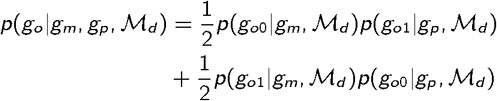
reflecting the uncertainty in parental origin of the offspring haplotypes, and where
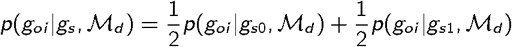
models uncertainty in which parental haplotype is inherited. 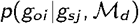 is the probability the haplotype *g_oi_* is inherited by the offspring given that the haplotype *g_sj_* is the one provided by the parent ***s*** for fertilization; it models *de novo* mutations.

For the female offspring X chromosome case we have the same form for 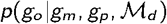 as the autosomal case, but
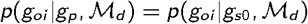

For the male offspring X chromosome case we have
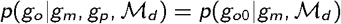

Finally, in the male offspring Y chromosome case we simply have
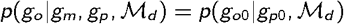

#### Cancer genotype prior model

The cancer genotype prior model, 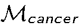, is used to assign probability to *cancer genotypes;* a pair of regular genotypes, *g_cancer_* = (*g_germ_, g_som_*), where *g_germ_* is the germline and *g_som_* is acquired somatically. The model must explain both the germline and the somatic genotypes. No assumptions of either germline or somatic genotype ploidy are made.

The model 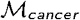 is the composition of two separate models: a germline prior model, 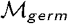 (e.g. the coalescent-HWE model); and a *conditional somatic model,* 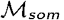. We then have (omitting models for brevity)
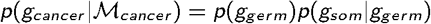

The second term *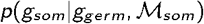* models the *pattern* of somatic haplotypes. In the simplest case when |*g_som_*| = 1 (i.e. there is a single somatic haplotype) then we just have
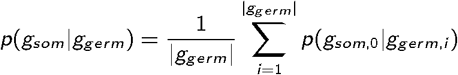
where 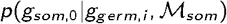 is the probability of observing the somatic haplotype given the germline haplotype *g_germ,i_* suffers some mutational event assigned by the somatic mutation model.

More interesting is when |*g_som_*| > 1 (i.e. there are more than one somatic haplotypes). In principle, we must consider that any of the somatic haplotypes could have originated from either the germline *or any other somatic haplotype;* we should consider possible tumour phylogenies. We have not implemented such a model in Octopus; we assume independence between somatic haplotypes:
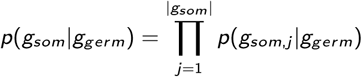

### Genotype posterior models

Genotype posterior models combine genotype prior models with genotype likelihood models. The three core genotype models currently implemented in Octopus **(Supplementary Fig. 10).**

#### Population genotype model

All samples have known ploidy and copy number, so the likelihood function of a genotype *g* given reads 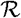 is
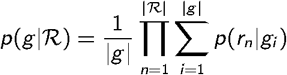
where |*g*| is the ploidy and 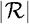 is the number of reads. The joint genotype posterior for *S* samples is therefore given by
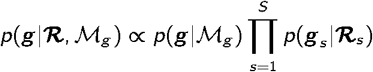
where the genotype prior model, 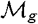, is either the uniform or HWE-coalescent prior. Unfortunately, the number of genotype combinations ***g*** grows exponentially in the number of samples ***S***, so we cannot evaluate the full posterior distribution in general. Therefore, other than for trivial cases, we first approximate 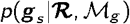 - the sample marginal genotype posterior distribution under the HWE model (without mutations), and use these marginal probabilities to select K genotype combinations ***g****_i_*, …, ***g****_K_* (K is user-defined) to evaluate under the full joint genotype model. The approximate posterior marginals are computed with expectation maximization (EM).

#### Individual genotype model

The individual model is simply a case of the population model, without the initial approximation step; this model is always fully evaluated.

#### Trio genotype model

The trio genotype model has the same likelihood function as the population model, but assigns prior probabilities to genotype combinations with the trio genotype prior model. Like for the population model, this model is intractable in general, so we only evaluate the posterior partially. Briefly, we evaluate approximate marginal probabilities for each sample under independence, and use these likelihoods to first combine parental genotypes, and then formulate a list of trio genotypes by combining parental combinations with offspring genotypes.

#### Subclone genotype model

Unlike the other genotype posterior models, this model does not assume known copy number - or mixture frequency - of sample genotypes. The model assumes all read observations originate from the same underlying genotype, but may have been observed at different mixture frequencies. The mixture frequencies for each sample therefore becomes a latent variable that we infer. We use Dirichlet distributions to assign probability to mixture frequencies, so different mixture priors may be specified for each sample by controlling the concentration parameter of the respective Dirichlet distribution. The joint posterior distribution is
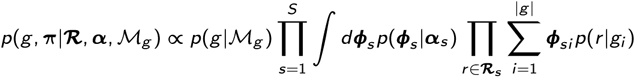
where ***α*** are prior concentration parameters, ***ϕ*** are mixture frequencies, and 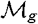 is the genotype prior model. We cannot compute this model exactly - or even partially - due to the integral over mixture frequencies. We therefore compute approximate posteriors for each latent variable using variational Bayes (VB). Namely, we assume the posterior distribution has factorization
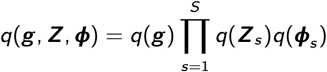
where ***Z****_s_* is a latent binary indicator matrix specifying read-component assignments. The associated probabilities, *q*(***Z_snk_***), are often called *responsibilities* - the responsibility haplotype *k* assumes for read *n* in sample *s.* With this factorisation, the posterior distributions for each latent variable is conjugate with the prior; we infer Dirichlet posterior distributions. An important feature of the VB approximation is that we can easily calculate a *lower bound* for the data likelihood - the model *evidence.*

### Calling models

Each calling model is responsible for calling variants given candidate alleles, haplotypes, and haplotype likelihoods. Although calling models are free to choose which latent variables and genotype models to use, all calling models must be able to infer posterior distributions over candidate haplotypes (for filtering), and genotypes (for phasing).

#### Individual

Unsurprisingly, the individual calling model uses the individual genotype model for genotype inference. Haplotype posteriors are computed by marginalising over the genotype posterior distribution:
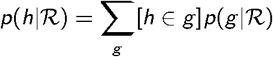
where *h* ∊ *g* is true if *h* occurs in *g* at least once. To call variants, we first calculate the posterior probability of all candidate non-reference alleles by marginalising over the genotype posterior distribution:
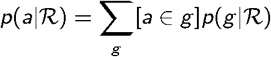
where *a* ∊ *g* is true if the any (*a* ∊ *h*) for *h* : *g* (i.e. if *a* occurs in any of the haplotypes in *g*). We then select alleles with posterior probability above some user-specified threshold.

Next, we identify the genotype with the greatest posterior probability (i.e. the MAP genotype). All selected alleles that appear on the MAP genotype are called, where the variant quality is determined by the marginalised allele posterior computed previously.

For each called allele, we then call genotypes at the loci of those alleles. In particular, for each allele we identify the alleles present in the called genotype at the loci of the allele, and once again marginalise over the genotype posterior distribution to compute the posterior probability for that local genotype. This is used for the genotype quality score.

#### Population

Inference for the population model is similar to the individual; variant calls are made based on sample marginal posteriors.

#### Trio

The trio calling model first infers a joint genotype posterior distribution 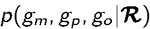 with the trio genotype posterior distribution. We then infer sample marginal genotype posteriors for each sample by marginalising over the joint posterior distribution. Haplotype posteriors are calculated by integrating over the marginal posterior:
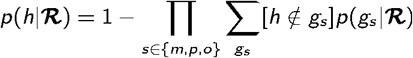

To calculate the probability an allele segregates in the trio, we also integrate over the joint genotype posterior distribution:
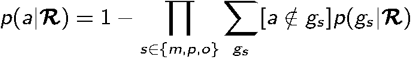

Similarly, we calculate the posterior probability an allele is *de novo* in the child:
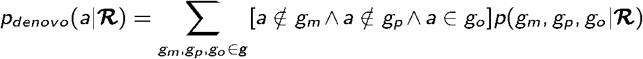

Finally, we find the MAP genotype combination using the joint genotype posterior distribution, and only call variants that are included in this trio.

#### Cancer

The cancer calling model is intended to detect somatic mutations in a set of tumour samples from a single individual. The set of samples may also include a sample that is not expected to contain somatic mutations - a normal or control sample. We attempt to model three data characteristics that result from tumour biology and experimental protocol:

i. There are no somatic mutational events; reads are generated from a clean germline.
ii. Copy number changes have occurred, but no somatic mutations.
iii. Somatic mutations have occurred, *and* possibly copy number changes.

Each of these three cases is modelled by fitting a unique genotype posterior model:

i. The individual model with any germline genotype prior model (all read observations are merged): 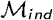.
ii. The subclone model with a *germline* genotype prior model (e.g. Coalescent-HWE): 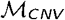.
iii. The subclone model with the *cancer* genotype prior model: 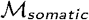.

The posterior probability for each model is calculated using Bayes theorem:
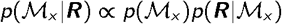
where 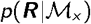 is the *evidence* for model ***x***.

For the somatic case, 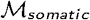, we must also infer the number of segregating somatic haplotypes. To do this we start by assuming a single somatic haplotype, and incrementally add more while the evidence for the model is the greatest observed so far, up to a user-defined limit.

For inference, we must marginalise over models. For example, we calculate posteriors for germline genotypes by marginalisation:
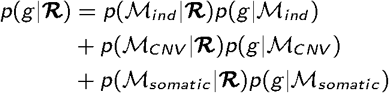
where 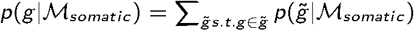 (i.e. marginalise over cancer genotypes that contain the matching germline component).

The posterior probability of an allele ***a*** segregating in the germline is then:
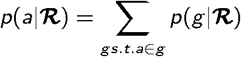

Germline candidates are called if the posterior is above a user-defined threshold, and if the candidate is present in the called germline genotype. If the allele is not called in the germline then it is added to a list of candidate somatic alleles.

To calculate the posterior probability an allele *a* is somatic, we marginalise over 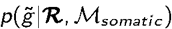, conditional on the somatic mutation frequency being above a user-defined threshold, *τ.*

First, we calculate the posterior mass for *credible* somatic frequencies. Supposing that we inferred a model with *K* somatic haplotypes, we assign probability to each somatic haplotype *k* = 1 … *K* if it occurs as a frequency above *τ,* 
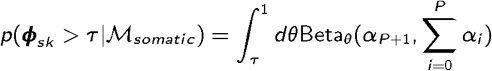
where the equality holds since the posterior distribution for ***ϕ_s_*** is Dirichlet. The overall credible somatic mass ***λ_s_*** is then calculated with
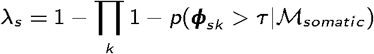

Finally we set ***λ*** = 1 – **∏ *λ_s_***. We then calculate the posterior probability that an allele is a somatic mutation by marginalisation:
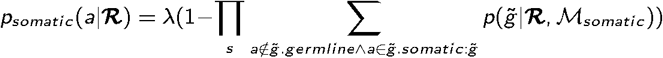

If this probability is greater than some user-defined threshold, we call the allele somatic.

For both called germline and somatic mutations, we also calculate the probability that the variant segregates regardless of classification, by marginalising over all three models.

#### Polyclone

The polyclone calling model is similar to the cancer calling model without the third somatic mutation model; It compares the individual genotype model with haploid genotypes, with the subclone genotype model, where the number haplotypes is determined iteratively by comparing model evidences. The genotype prior model in both cases is either the uniform of Coalescent-HWE model (depending on user choice).

### Probabilistic phasing

Although each caller implements different genotype models, each is required to infer the *marginal* posterior probability of genotypes for each sample. This posterior distribution is used to infer physical phasing of called variant sites. Direct read data is not required as all information available from the reads, in addition to any prior information, is already contained in the posterior distribution. The advantage of this approach is most evident when calling trios as the genotype prior can be strongly informative about phase due to identity by decent. The method applies to genotypes of arbitrary zygosity and is therefore applicable to non-diploid samples.

Samples are phased independently; the marginal genotype posterior distribution for each sample is used for phasing. First, all genotypes in the domain of the posterior distribution are partitioned into *phase complement* sets. All genotypes in a phase complement set share exactly the same alleles, although they may appear on different haplotypes. There is only one such partitioning possible for any set of genotypes. We calculate the entropy of each set with respect to the normalised posterior probabilities of the genotypes contained in the set. A low entropy implies little uncertainty in the phase of the alleles present in the set. To account for uncertainty in the samples genotype, we marginalise over all phase complement sets by taking the weighted sum of each sets entropy, where the weight for each set is the sum of the unnormalised posterior probabilities. We termed this weighted average a *phase score,* 
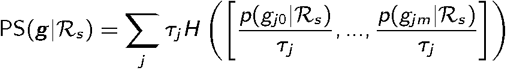
where 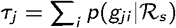 is the sum of genotype posteriors in phase complement set *j,* and *H* is the binary Shannon entropy function.

Lower phase scores indicate less ambiguous phasing, even if there is high uncertainty in the samples genotype. If the phase score for a set of genotypes is lower than a user-defined threshold the region is considered phased, otherwise, each genotype in the set can be broken into two parts which results in two unphased sets of partial genotypes. The phase scores of these two partial sets is never greater than the phase score of the original genotype set. The phasing algorithm therefore iteratively finds the smallest set of breakpoints such that the partial genotype sets defined by those breakpoints are all phased.

### Variant filtering

As with any model, there are some error modes that are not well captured by Octopus calling models which can lead to false inferences. For example, Octopus assumes read sequencing and mapping errors are independent which is not true in general. To identify false calls due to model error, we developed classifiers to filter Octopuss raw calls using statistics, called *measures* in Octopus, that may be derived directly from the input read data.

#### Hard filtering

Hard filters are Boolean expressions where the terms of the expression are comparison operations. Currently, only *or* binary operations are permitted; if any of the individual operations is true then the call is filtered. Different filter expressions can be specified for germline variant, *de novo,* somatic, and homozygous reference calls.

#### Random forest filtering

Octopus uses the Ranger library ^46^ for random forest classification. Different random forests may be used for germline and somatic calls.

### BAM realignments

Octopus is capable of producing realigned BAM files that provide further evidence of a calls reliability. These BAM files are especially helpful in cases where there are complex indel variants and the input alignments are significantly different from the alignments supported by the called haplotypes. The realignment process for each read is:

1. Identify the called haplotype where the read originated from.
2. Align the haplotype to the reference sequence.
3. Align the read to the called generating haplotype (mismatches due to sequencing or calling errors).
4. Merge the two alignments to obtain a single alignment to the reference.

For the first step, the called haplotype with the maximum likelihood of generating the read is used. If there are more than one called haplotypes that have equal likelihood, then we label the read *ambiguous.* For realignment, ambiguous reads are assigned randomly to one of the equally well supported haplotypes. The second step is trivial in Octopus since haplotypes are defined explicitly by alleles that are reported in the VCF output. The third step is computed using the Viterbi alignment found from calculating the maximum likelihood in the first step.

Since a hard choice of generating haplotype is made in the first step, we can report this information by generating separate BAM files for each called haplotype, and another for ambiguous reads.

### Synthetic tumours

To generate synthetic tumour BAM files we followed a similar approach to Ewing et al. ^32^ with important differences. First, we obtained unmapped reads from the GIAB’s NA12878 high coverage HiSeq 2500 set (~ 300x total). From the full high coverage set, we extracted four non-overlapping subsets such that the average depths in the four subsets would be 30x, 35x, 60x, and 65x. To maintain realistic sequecing conditions, we ensured that reads from same library and sequenced on the same lane were kept together. The 30x and 35x read sets were used as the normal samples, while the 60x and 65x read sets were used to make the synthetic tumour samples.

For each of the two neo-synthetic-tumour BAMs, we then performed a haplotype-based read assignment and realignment using Octopus’ ‘split’ BAM realignment feature. This results in three BAM files; two containing reads that were assigned to a called germline haplotype, and another containing ambiguous reads. The purpose of the assignment step is to ensure spike in mutations fall on the same germline haplotype. The realignment step is to ensure consistency of spike-in location. Neither of which are guaranteed by the method described in Ewing et al. due to limitations with BAM-Surgeon. In summary, this procedure results in three ‘cleaned’ BAM subsets, each of which should be haploid and contain few alignment errors.

We then simulated two sets of somatic mutations by sampling putative real somatic mutations called by the PCAWG consortium. In particular, we uniformly sampled PCAWG calls from skin tumours and from breast tumour, in order to achieve mutation densities close to 1Kb^−1^ and 1Mb^−1^ for the skin and breast sets, respectively. These densities were chosen from the upper expected range for each tumour type ^29^. For each sampled mutation, we uniformly assigned a spike-in variant allele frequency from 0.025 to 0.5 (from 20 equally spaced bins) for breast mutations, and 0.025 to 0.2 (from 20 equally spaced bins) for breast mutations. Although this frequency distribution is unlikely to be biologically realistic, we used this approach to evaluate the performance of each algorithm across a range of variant allele frequencies, and to the best of our knowledge, none of the evaluated methods consider genome-wide sub-clonal tumour architecture when calling variants. Each sampled mutation was finally assigned to a random germline haplotype.

We used a slightly modified version of BAMSurgeon (https://github.com/dancooke/bamsurgeon) to spike in mutations into each putative haploid neo-tumour BAM. The main modification that we made to BAMSurgeon was to ensure any ‘pad’ sequence used for deletion spike-ins came from the originating germline haplotype, rather than the reference sequence. We also needed to make minor modifications to handle our split BAM files, which may not contain proper read pairs. The spike-in VAF used for the BAM containing ambiguous reads, which should contain reads from both parental haplotypes, was always half the chosen spike-in VAF.

Finally, we created paired raw synthetic tumour FASTQ files by merging and extracting reads from the three spiked haploid BAM files. We emphasise that this final step removes all alignment and phasing information.

### Code availability

Octopus source code and documentation is freely available under the MIT licence from https://github.com/luntergroup/octopus. Custom code used for data analysis is available from monospace https://github.com/luntergroup/octopus-paper.

### Data availability

All germline data used in this manuscript is publicly available from Genome in a Bottle, Precision FDA, and EGA. The WGS500 trio data used for *de novo* analysis is not publicly available. The synthetic tumour data is freely available from https://storage.googleapis.com/luntergroup/syntumour.

### Author contributions

D.C and G.L designed the algorithm and wrote the manuscript. D.C implemented the algorithm and performed the the evaluation. D.W provided data for the synthetic tumours and critically reviewed the manuscript.

